## Supplementary Information for "The co-receptor Tetraspanin12 directly captures Norrin to promote ligand-specific β-catenin signaling"

### Table S1. List of measured affinities and kinetic constants

| Bait (biosensor) | Analyte (in solution) | Steady- state $K_D$ (nM) | Kinetic $K_D$ (nM) | $K_{on}$ ($nM^{-1}$ $s^{-1}$) x $10^{-4}$ | $K_{off}$ ($s^{-1}$) x $10^{-4}$ | Relating to figure | Notes |
|---|---|---|---|---|---|---|---|
| Tspan12ΔC | Norrin | 10.4 ± 1.2 | 7.4 ± 1.4 | 1.9 ± 0.3 | 14 ± 2 | 1A-D, 4A | $h$ = 0.98 ± 0.1 |
| MBP-Norrin | Tspan12 LEL | 17.9 ± 3.0 | 16 ± 3 | 0.70 ± 0.08 | 11 ± 2 | 1E | |
| Tspan12-LEL11 | Norrin | | n.b. at 100 nM | | | 1F | |
| MBP-Norrin | Tspan12ΔC in nanodiscs | 18.0 ± 4.0 | 34 ± 11 | 1.0 ± 0.3 | 34 ± 3 | S2A | |
| MBP-Norrin | Tspan12ΔC in GDN | | n.d. ( > 100 nM) | | | S2B | |
| Fzd4 | Tspan12 LEL (MBP-tagged) | | n.b. at 18 µM | | | S2C | |
| Tspan12ΔC | Fzd4 CRDL | | n.b. at 32 µM | | | S2D | |
| MBP-Norrin | Fzd4 CRDL | 122 ± 38 | 340 ± 42 | 0.43 ± 0.05 | 150 ± 10 | S2E | |
| Tspan12ΔC | WT Norrin | | 2.9 ± 0.7 | 1.9 ± 0.5 | 4.8 ± 0.5 | | Measured at 32 nM Norrin |
| WT Tspan12 | WT Norrin | | 2.5 ± 0.2 | 1.1 ± 0.1 | 2.7 ± 0.2 | 2, S5 | n = 6; Measured at 32 nM Norrin |
| WT Tspan12 | Norrin R107E/R115E | | 210 ± 24 | 1.9 ± 0.5 | 520 ± 30 | 2E, S5A/C/E | Measured at 32 nM Norrin |
| WT Tspan12 | Norrin T117Y/T119Y | | 30 ± 7 | 0.32 ± 0.02 | 9.6 ± 1.9 | 2E, S5A/C/E | Measured at 32 nM Norrin |
| WT Tspan12 | Norrin K102E/R121E | | 240 ± 28 | 1.7 ± 0.3 | 380 ± 40 | 2E, S5A/C/E | Measured at 32 nM Norrin |
| WT Tspan12 | Norrin S82D | | 5.4 ± 0.4 | 1.5 ± 0.2 | 8.4 ± 1.5 | 2E, S5A/C/E | Measured at 32 nM Norrin |
| Tspan12 E173K/D175K | WT Norrin | | 8.5 ± 1.2 | 0.75 ± 0.07 | 6.3 ± 0.3 | 2F, S5B/D/F | Measured at 32 nM Norrin |
| Tspan12 L198Y | WT Norrin | | 1.9 ± 0.16 | 0.95 ± 0.05 | 1.7 ± 0.1 | 2F, S5B/D/F | Measured at 32 nM Norrin |
| Tspan12 E196K/S199K | WT Norrin | | 14 ± 1.1 | 0.70 ± 0.09 | 9.5 ± 0.4 | 2F, S5B/D/F | Measured at 32 nM Norrin |
| Tspan12 E170K | WT Norrin | | 16 ± 1.3 | 0.66 ± 0.03 | 10 ± 0.6 | 2F, S5B/D/F | Measured at 32 nM Norrin |
| Tspan12 E173K/D175K | Norrin R107E/R115E | | n.b. at 32 nM | | | S6 | Measured at 32 nM Norrin |
| Tspan12 L198Y | Norrin T117Y/T119Y | | 23 ± 4.6 | 0.34 ± 0.09 | 8.7 ± 4.0 | S6 | Measured at 32 nM Norrin |
| Tspan12 E196K/S199K | Norrin K102E/R121E | | 1.3 ± 0.06 | 6.7 ± 0.3 | 8.8 ± 0.2 | S6 | Measured at 32 nM Norrin |

| | | | | | | | |
|---|---|---|---|---|---|---|---|
| Tspan12 E170K | Norrin S82D | | 5.2 ± 0.7 | 1.5 ± 0.03 | 7.7 ± 1.2 | S6 | Measured at 32 nM Norrin |
| Fzd4 | Norrin dimer | 3.29 ± 0.17 | n.d. | 3.1 ± 0.4 | 10 ± 1 [a] | 4A, S11A | *h* = 2.8 ± 0.6 |
| Tspan12/Fzd4 dimer | Norrin dimer | 2.18 ± 0.10 | n.d. | 4.3 ± 0.6 | 9.5 ± 1.0 [a] | 4A, S11B | *h* = 1.7 ± 0.1 |
| Tspan12ΔC | Norrin monomer | 820 ± 100 | 80 ± 35 | 0.52 ± 0.20 | 41 ± 9 | 4B | *h* = 1.0 ± 0.1 |
| Fzd4 | Norrin monomer | 13.2 ± 1.2 | n.d. | 1.9 ± 0.3 | 26 ± 4 [a] | 4B, S11C | *h* = 2.0 ± 0.3 |
| Tspan12/Fzd4 dimer | Norrin monomer | 11.5 ± 1.5 | n.d. | 1.9 ± 0.3 | 22 ± 4 [a] | 4B, S11D | *h* = 1.0 ± 0.1 |
| Fzd4[b] | Dvl2 DEP | 183 ± 24 | 220 ± 28 | 0.62 ± 0.07 | 140 ± 7 | 5B-C | |
| Tspan12/Fzd4 dimer [b] | Dvl2 DEP | 279 ± 46 | 280 ± 49 | 0.45 ± 0.07 | 130 ± 6 | 5C | |
| Fzd4 [b] + Norrin | Dvl2 DEP | 161 ± 21 | 160 ± 13 | 0.82 ± 0.06 | 130 ± 4 | 5D | |
| Tspan12/Fzd4 dimer [b] + Norrin | Dvl2 DEP | 274 ± 39 | 280 ± 110 | 0.43 ± 0.17 | 120 ± 6 | 5D | |

Note: Values represent mean ± S.E.M. for n=3 independent experiments unless otherwise noted.
Steady state $K_D$ was determined for experiments in which > 3 different concentrations of analyte were measured.
Bait receptors (Tspan12, Fzd4, and Tspan12/Fzd4) were embedded in MSP1D1 nanodiscs (48:32:20 POPC:POPG:Cholesterol) unless otherwise noted.
Abbreviations: n.b. = no binding. n.d. = not determined, i.e. fit was of low confidence.; *h* = hill coefficient.
[a] $K_{off}$ determined by steady state $K_D$*$K_{on}$, not by direct fitting of dissociation traces due to fits of low confidence.
[b] For Dvl2 DEP binding, the bait receptor was inserted into MSP1E3D1 nanodiscs (70:5:20 POPC:PI(4,5)$P_2$:Cholesterol)

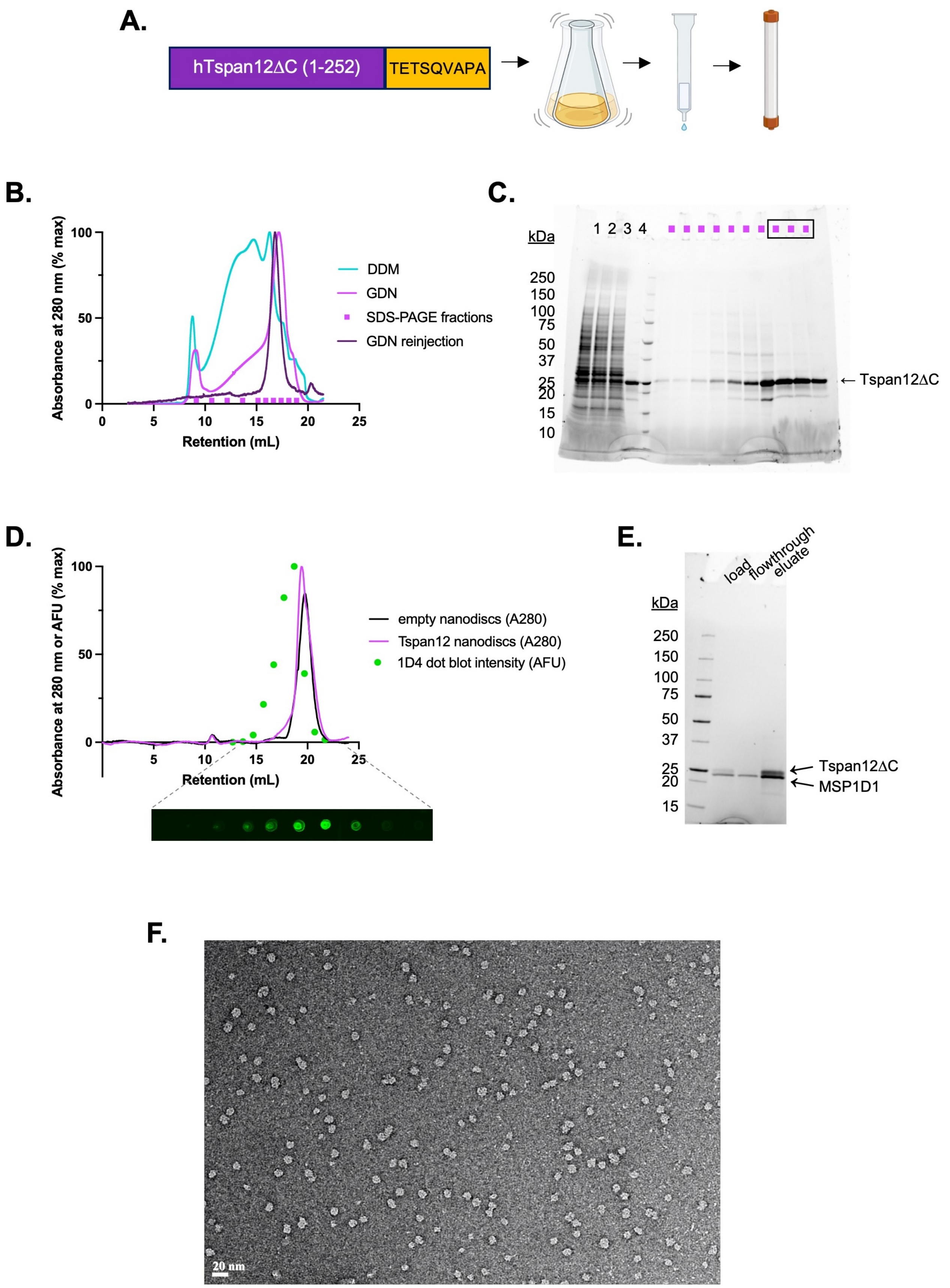


**Figure S1. Tspan12 purification. A.** Schematic of Tspan12 purification. Human Tspan12ΔC (truncated at residue 252 after the fourth transmembrane domain), was tagged with the Rho1D4 tag TETSQVAPA and expressed in SF9 cells by baculovirus. Detergent-solubilized Tspan12 was affinity purified on anti-Rho1D4 antibody resin followed by size exclusion chromatography. Figure created with BioRender (BioRender.com/q49x347). **B**. Superose 6 10/300 size exclusion traces of Tspan12 purified in DDM (teal) or exchanged into GDN during the affinity chromatography step (pink); the final concentrated product in GDN showed no signs of aggregation (purple). Indicated fractions of GDN-solubilized Tspan12 (pink trace) were run on **C.** SDS-PAGE; boxed fractions were pooled and concentrated. **D.** Size exclusion traces of empty nanodiscs (black) or Tspan12 reconstituted into excess MSP1D1 nanodiscs (pink). Tspan12 content of fractions was quantified by dot blot (anti-Rho1D4) and dot intensity was plotted accordingly (green). **E.** Peak Tspan12-containing fractions from (D) were affinity purified on anti-Rho1D4 resin. Shown is an SDS-PAGE gel of the load, flowthrough, and eluate. **F.** Uranyl acetate negative stain micrograph of final nanodisc-reconstituted Tspan12. Scale bar is 20 nm.

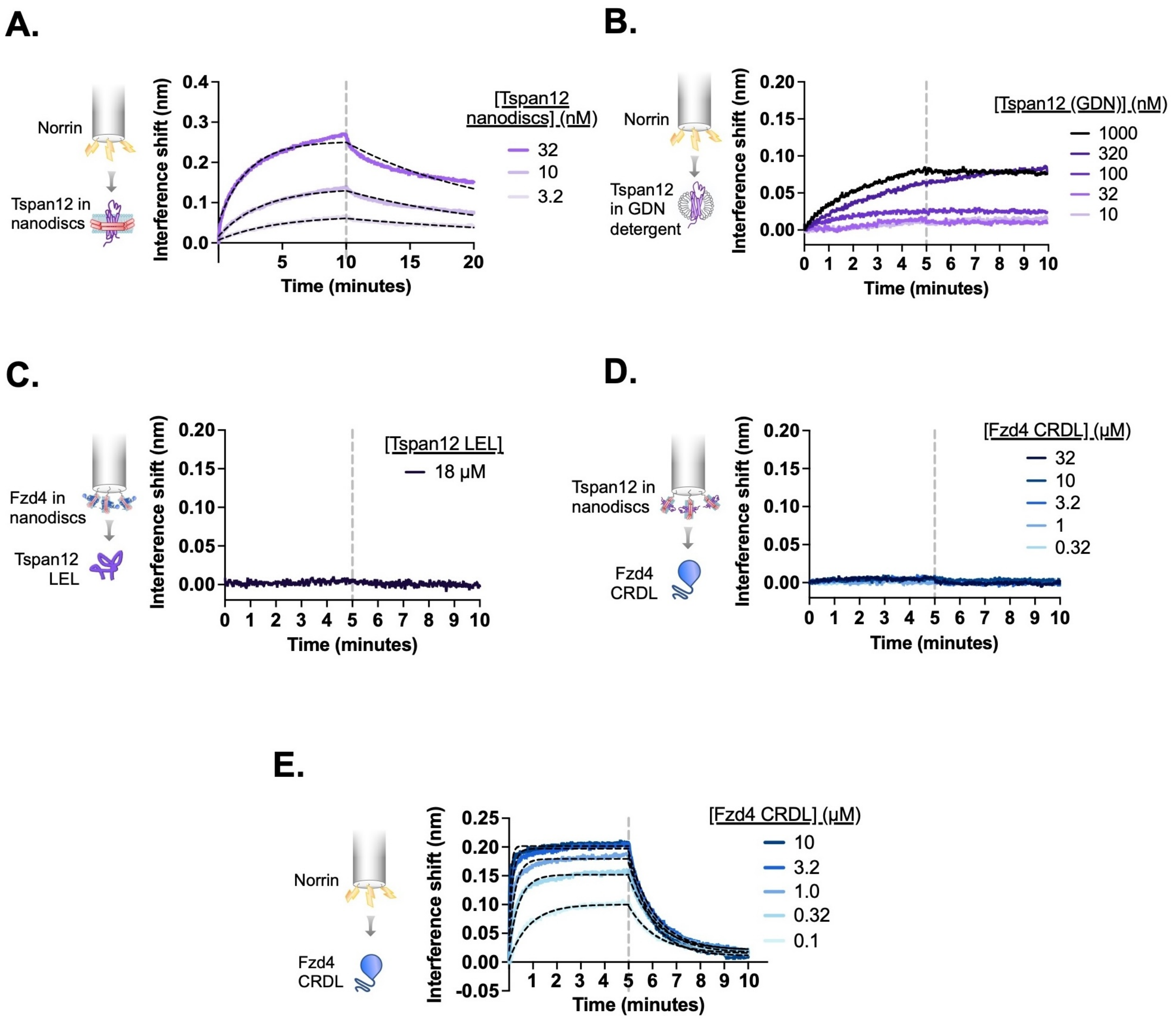


**Figure S2. Tspan12 and Fzd4 each bind Norrin, but not one another, with high affinity through their extracellular domains. A.** BLI traces showing non-biotinylated Tspan12 nanodiscs in solution binding to biosensor-immobilized biotinylated MBP-Norrin (flipped setup compared to Fig 1A-B). Overlaid fits (black dashed line) give an apparent affinity of 34 ± 11 nM. **B.** BLI traces showing weak binding of indicated concentrations of Tspan12 in GDN detergent to biosensor-immobilized biotinylated MBP-Norrin. **C.** BLI trace showing no binding of MBP-tagged Tspan12 LEL at 18 $\mu$M to biosensors loaded with nanodisc-embedded Fzd4. **D.** BLI traces showing no binding of the soluble Fzd4 CRDL up to 32 $\mu$M to biosensors loaded with nanodisc-embedded Tspan12. **E.** BLI traces showing binding of the Fzd4 CRDL to biosensor-immobilized MBP-Norrin, which reaches saturation at 3.2 $\mu$M CRDL. Equilibrium binding values obtained from the overlaid fits (black dashed lines) give a steady-state affinity of 122 ± 38 nM (mean ± S.E.M.), in agreement with previously reported affinity measurements (Bang et al., 2018).

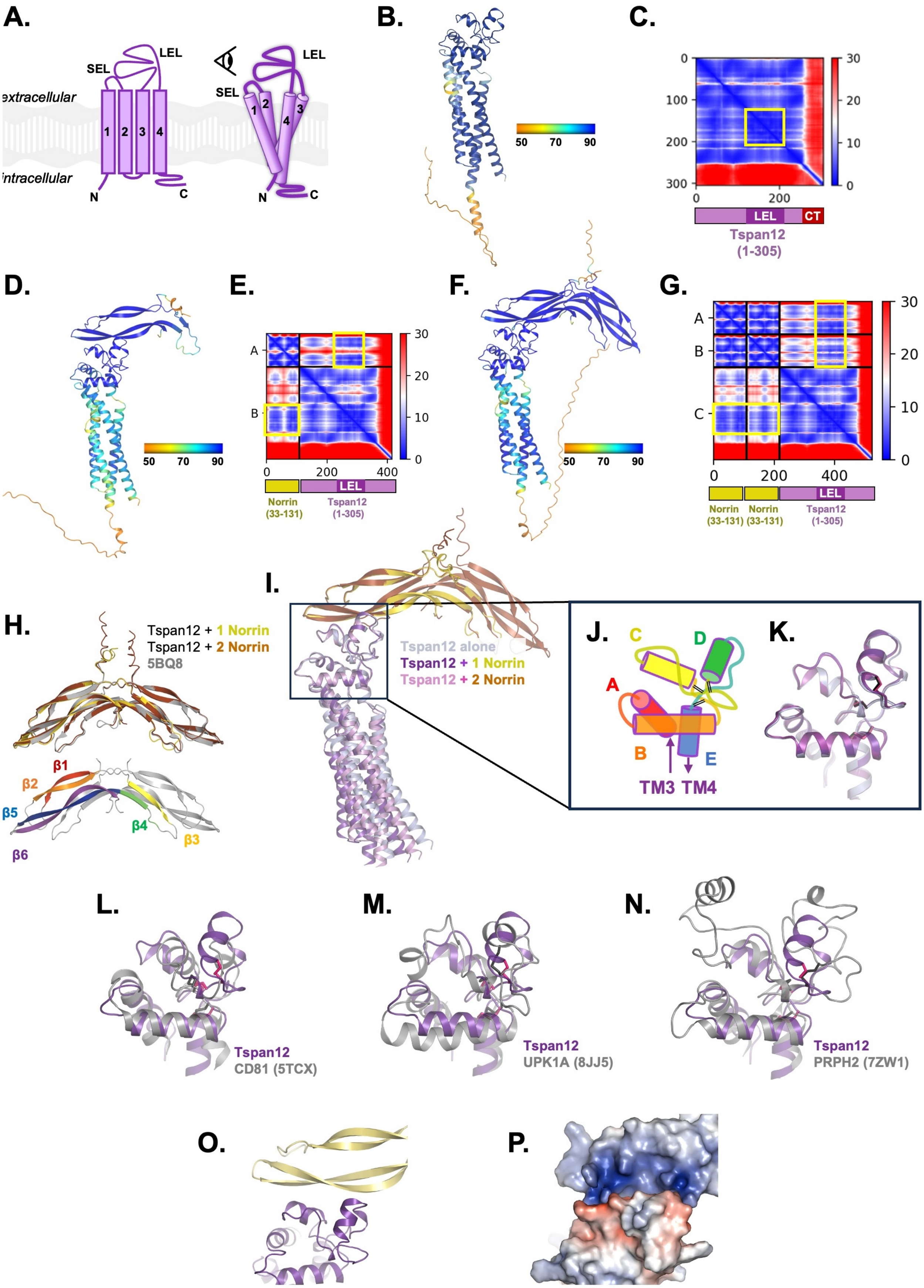

A.
SEL
LEL
extracellular
intracellular
1 2 3 4
N
C
SEL
LEL
1 2 4 3
N
C
B.
50 70 90
C.
0
100
200
300
0 200
30
20
10
0
LEL
CT
Tspan12
(1-305)
D.
50 70 90
E.
A
B
0 200 400
30
20
10
0
LEL
Norrin
(33-131)
Tspan12
(1-305)
F.
50 70 90
G.
A
B
C
0 200 400
30
20
10
0
LEL
Norrin
(33-131)
Norrin
(33-131)
Tspan12
(1-305)
H.
Tspan12 + 1 Norrin
Tspan12 + 2 Norrin
5BQ8
β1
β2
β5
β6
β4
β3
I.
Tspan12 alone
Tspan12 + 1 Norrin
Tspan12 + 2 Norrin
J.
C
D
A
B
E
TM3 TM4
K.
L.
Tspan12
CD81 (5TCX)
M.
Tspan12
UPK1A (8JJ5)
N.
Tspan12
PRPH2 (7ZW1)
O.
P.

**Figure S3. AlphaFold structural prediction of Tspan12 bound to Norrin. A.** Cartoon of tetraspanin structure, comprised of transmembrane helices 1-4, a small extracellular loop (SEL, between helices 1 and 2) and a large extracellular loop (LEL, between helices 3 and 4). Eye icon indicates the viewing angle of AlphaFold models in the remainder of this figure (B, D, F, I-P) relative to this cartoon. **B.** The structure of full-length human Tspan12 alone was predicted with AlphaFold. The best-scoring model is shown, colored by the per-residue predicted local distance difference test (pLDDT) confidence metric. **C.** The predicted aligned error (PAE) for the model in B. The position of the LEL (darker purple) appears as a darker blue square in the heat map with low PAE (yellow box). The position of C-terminal residues ("CT", red) is indicated; the heat map shows high PAE values for the C-terminus relative to the rest of the protein, indicating poor prediction of the relative positioning of the C-terminus. **D.** The structure of full-length human Tspan12 together with one Norrin protomer (residues 25-133) was predicted with AlphaFold-Multimer and the best-scoring model is shown, colored by pLDDT. **E.** The predicted aligned error for the model shown in D. The position of Norrin, Tspan12 and the LEL along the axes are indicated. Note low PAE between the LEL and parts of Norrin (yellow boxes). **F.** The structure of full-length human Tspan12 together with two copies of Norrin (residues 25-133) was predicted with AlphaFold-Multimer and the best-scoring model is shown, colored by pLDDT. **G.** The predicted aligned error for the model shown in F. The position of both copies of Norrin, Tspan12 and the LEL along the axes are indicated. Note low PAE between the LEL and parts of both Norrin protomers (yellow boxes). **H.** Top: The structure of Norrin within the predicted structure of Tspan12 + 1 Norrin protomer (yellow), and within the predicted structure of Tspan12 + 2 Norrin protomers (orange), matches the crystal structure of Norrin (5BQ8 chains A and B; gray) with RMSDs of 0.383, and 0.413 Å respectively. Below: β strands 1, 2, 3, 4, 5 and 6 of one Norrin protomer, colored red, orange, yellow, green, blue and purple, respectively; strands 5 and 6 are predicted to comprise the Tspan12 binding site. **I.** The predicted model of Tspan12 alone (light blue) is similar to the predicted models with one (purple/yellow) or two (pink/orange) copies of Norrin, which each align with RMSDs of 1.247 and 1.057 Å, respectively, to the model of Tspan12 alone. Aligning only the LELs as shown gives RMSDs of 0.293 and 0.341 Å, respectively, and illustrates the slight variation in the predicted angle between the TMs and the LEL. The predicted position and orientation of Norrin relative to the LEL is unchanged between the one-protomer and two-protomer models. **J.** Tetraspanin LELs are composed of helices A, B, C, D, and E. Helices C and D are the least conserved and are implicated in binding partner interactions (Susa et al., 2023). Black double lines represent disulfide bridges. Viewing angle is indicated by the eye in A. **K.** Close up of aligned LELs from figure I. **L.** The predicted structure of the Tspan12 LEL (from the 1 Tspan12 : 1 Norrin protomer model; residues 123-215; purple) aligns to the experimentally-determined structure of CD81 LEL (5TCX residues 123-198; (gray), with an RMSD of 3.997 Å. CD81 is a C4 tetraspanin; disulfide bonds for Tspan12 and CD81 are shown in pink and black, respectively. **M.** The predicted Tspan12 LEL (purple) aligns to the Uroplakin 1A LEL (8JJ5, residues 125-226; gray), a C6 tetraspanin, with an RMSD of 4.215 Å. **N.** The predicted Tspan12 LEL (purple) aligns to the Peripherin-2 LEL (7ZW1, residues 126-258; gray), a C6 tetraspanin, with an RMSD of 2.528 Å. **O.** The predicted interaction between Norrin and Tspan12 involves helices C and D of the LEL and residues on β5 and β6 of Norrin. **P**. The predicted interaction between Norrin and Tspan12 colored by surface electrostatics (APBS) shows a highly polar interaction involving a basic patch on Norrin and an acidic patch on Tspan12.

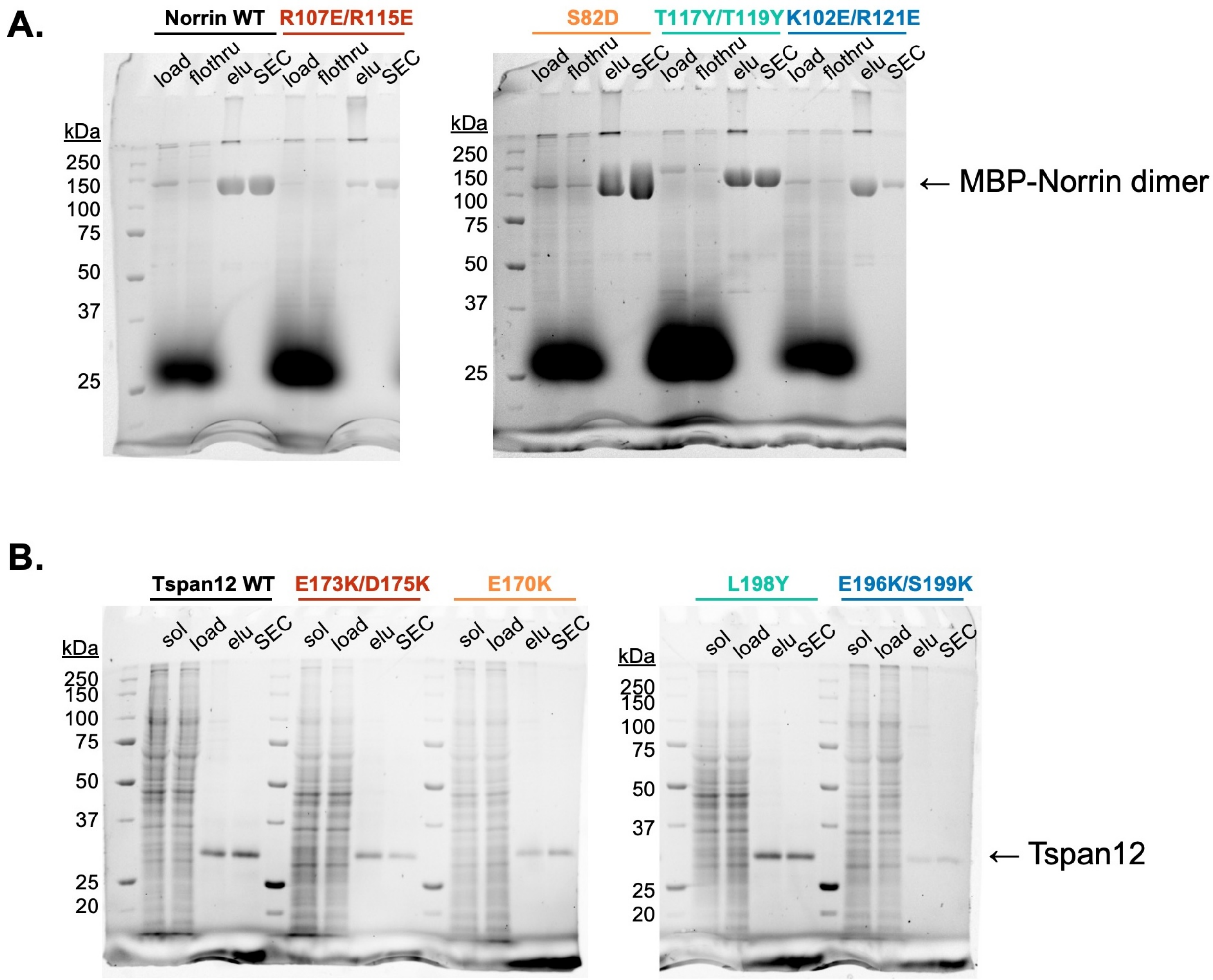


**Figure S4. Purification of Norrin and Tspan12 mutants. A.** Stain-free non-reducing SDS-PAGE gels of Sf9 supernatant media ("load"), amylose resin flowthrough ("flothru"), amylose eluate ("elu"), and pooled fractions after size exclusion ("SEC") for WT and indicated mutant MBP-Norrin. **B.** Stain-free SDS-PAGE gels of DDM-solubilized Expi293 membranes ("sol"), supernatant post-ultracentrifugation ("load"), 1D4 antibody resin eluate ("elu"), and pooled fractions after size exclusion ("SEC") for full-length WT and indicated mutant Tspan12.

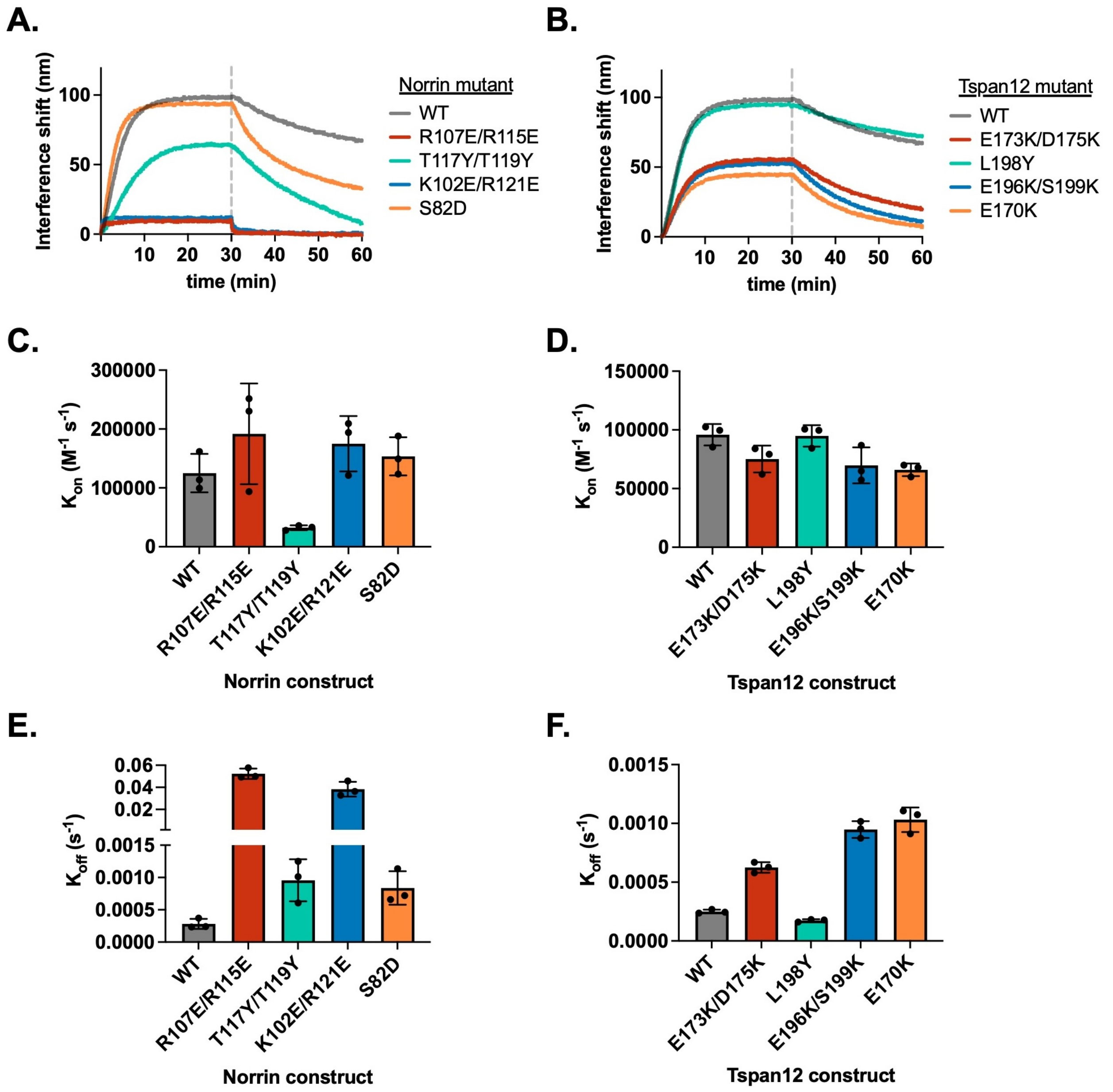


**Figure S5. Binding kinetics of Norrin and Tspan12 mutants. A.** Representative BLI association and dissociation traces of 32 nM WT or mutant Norrin binding to immobilized WT Tspan12, and for **B.** 32 nM WT Norrin binding to immobilized WT or mutant Tspan12. Colors correspond to sites within the binding interface defined in Fig. 2. **C.** and **D.** Rate constant $K_{on}$ (mean ± S.D.) calculated from association and dissociation fits to traces shown in A and B, respectively; each mutant was measured in triplicate. **E.** and **F.** Rate constant $K_{off}$ (mean ± S.D.) calculated from dissociation fits to traces shown in A and B, respectively; each mutant was measured in triplicate. Across the board, differences in $K_D$ for mutants compared to wildtype (Fig 2E and 2F) were driven largely by differences in $K_{off.}$ Kinetic constants reported in table S1.

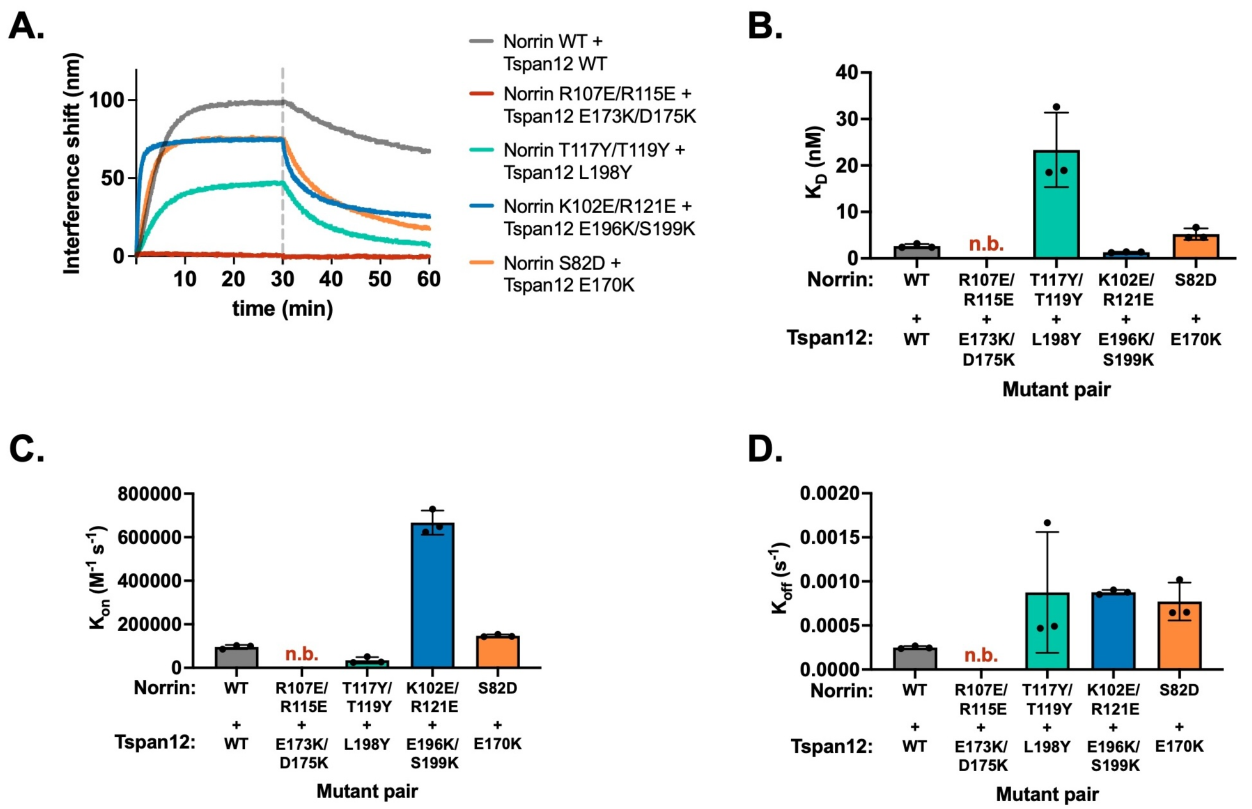


**Figure S6. Binding affinity and kinetics of mutant Norrin binding to mutant Tspan12. A.** Representative BLI association and dissociation traces showing 32 nM mutant Norrin binding to immobilized mutant Tspan12 relative to the WT/WT binding (grey) for mutants at Site 1 (red), Site 2 (teal), Site 3 (blue), and Site 4 (orange). For each site, mutated residues on Norrin and Tspan12 are predicted to interact according to the AlphaFold structure; in the case of charge-swapped mutations (i.e. Sites 1, 3, and 4), we hypothesized that the mutations to the two proteins may be compensatory. **B.** Binding affinities calculated from kinetic fits to traces in A. for the indicated mutant pairs. The charge-swapped mutants at Site 1, Norrin R107E/R115E and Tspan12 E173K/D175K, do not bind appreciably at 32 nM Norrin. At Site 3, charge-swapped Norrin K102E/R121E and Tspan12 196K/S199K binding is rescued to WT/WT levels compared to the much weaker binding affinities of either mutant for its wildtype counterpart: i.e., the mutations are compensatory. At Site 4, the Norrin S82D mutation partially rescues the deleterious effects of Tspan12 E170K. Data represent mean ± S.D. from three replicates. Binding affinities and kinetic constants reported in Table S1. **C.** $K_{on}$ and **D.** $K_{off}$, calculated from each individual trace shown in A, measured in triplicate. Plots represent three replicates ± S.D, and kinetic constants are reported in table S1.

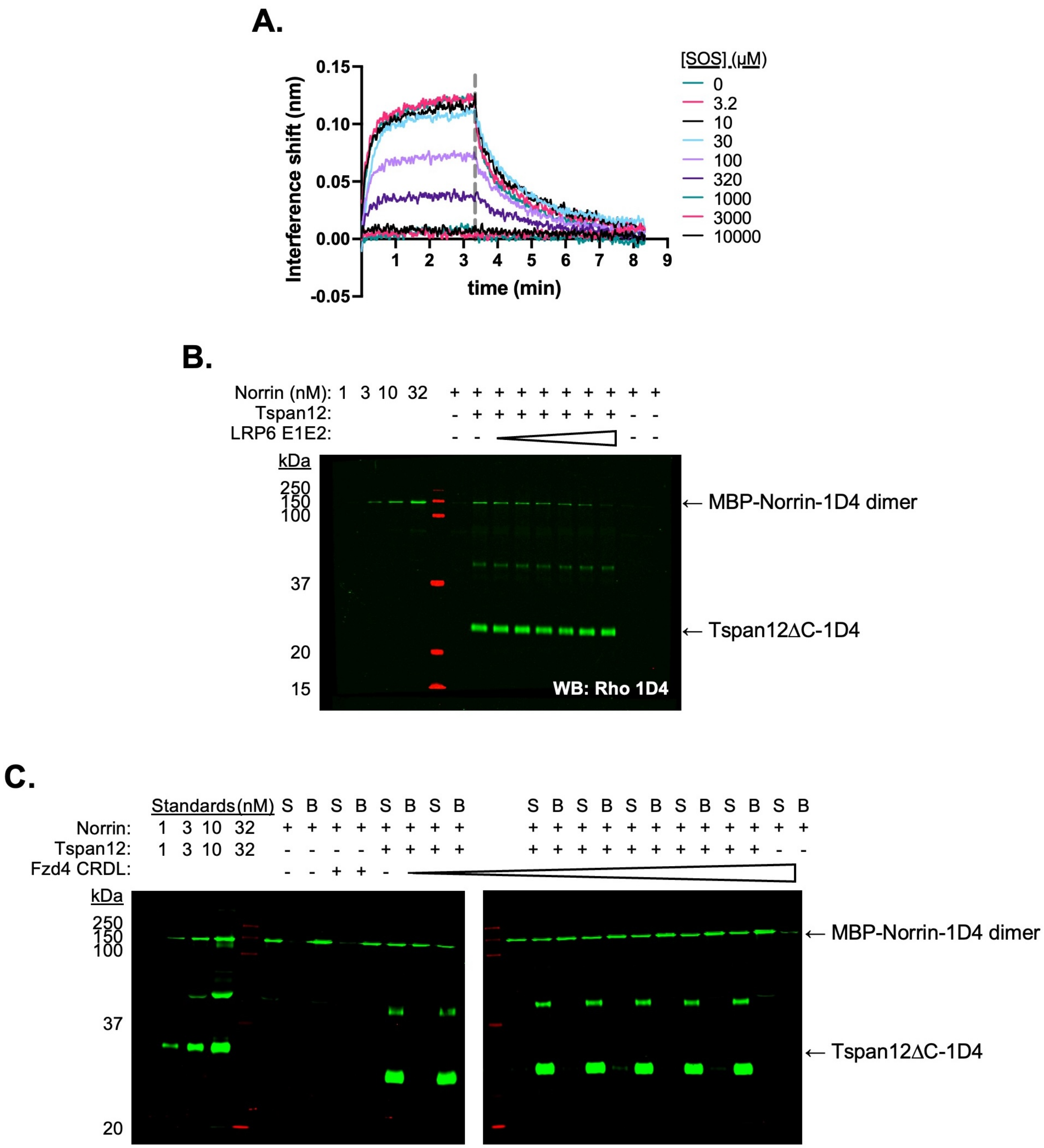


**Figure S7. Tspan12 Norrin binding can be competed with SOS and the LRP6 E1E2 domain but not the Fzd4 CRDL domain. A.** Representative BLI traces of 32 nM Norrin binding to biosensor-immobilized Tspan12 in the presence of increasing concentrations of SOS. **B.** Representative western blot of Norrin bound to Tspan12 or empty nanodiscs immobilized on paramagnetic particles in the presence of 0, 0.1, 0.4, 1.6, 6, 25, or 100 $\mu$M of purified LRP6 E1E2 domain, with both Norrin and Tspan12 detected by anti-Rho1D4 antibody. Norrin is detected within the dynamic range, as shown by concentration standards loaded to the left of the ladder. **C.** Representative western blot of Norrin in the supernatant (S) or remaining bound (B) to Tspan12 or empty nanodiscs immobilized on paramagnetic particles in the presence of 0, 0.1, 0.4, 1.6, 6, 25, or 100 $\mu$M of purified Fzd4 CRD-linker domain, with both Norrin and Tspan12 detected by anti-Rho1D4 antibody. Norrin and Tspan12 are detected within the dynamic range, as shown by concentration standards loaded at left. Tspan12 forms SDS-induced dimers, which can be detected at about 50 kDa.

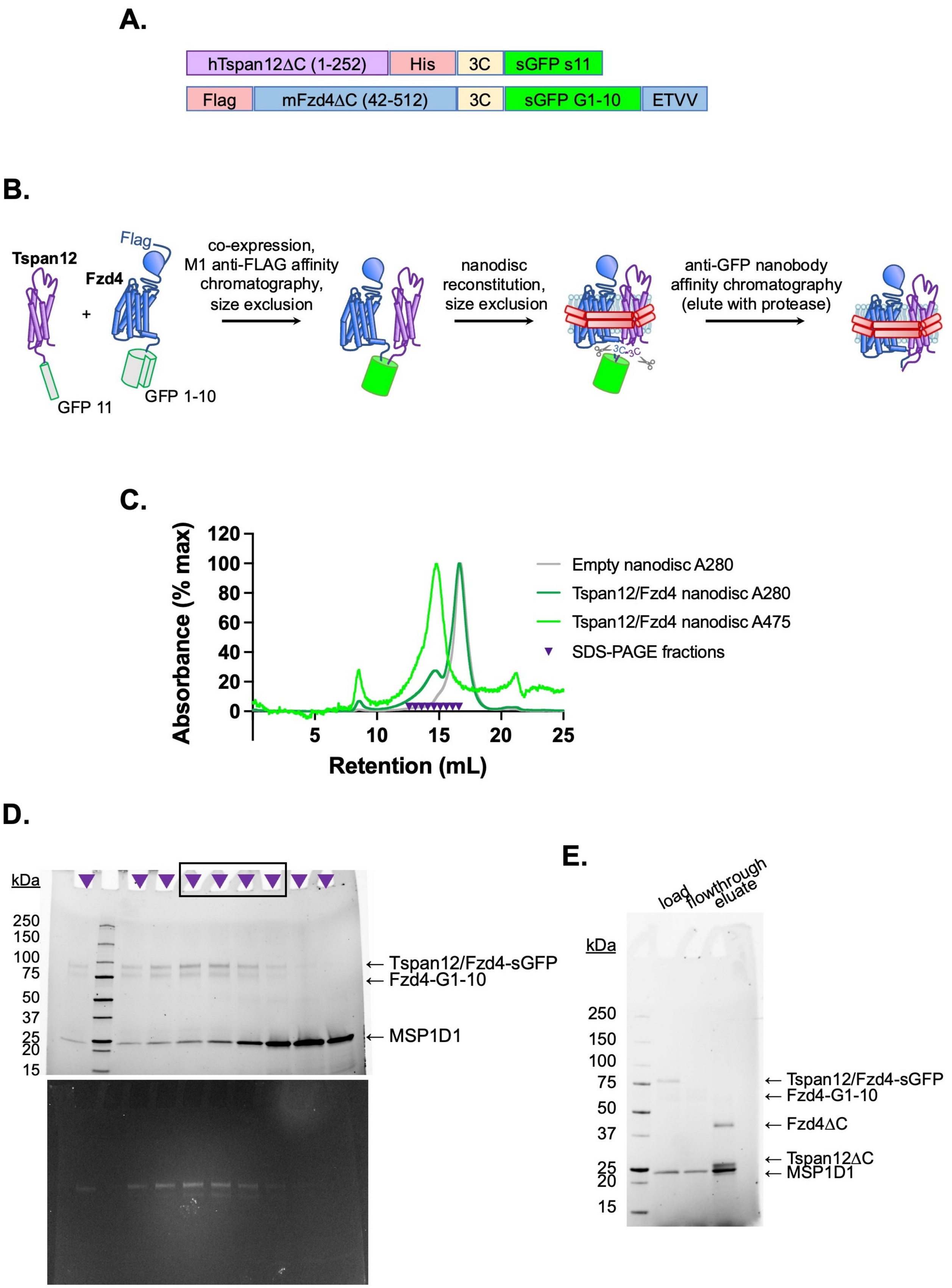


**Figure S8. Purification of Fzd4/Tspan12 dimer and insertion into nanodiscs. A.** Construct design of Tspan12 and Fzd4 C-terminally tagged with split GFP fragments s11 and s1-10, respectively, both downstream of a 3C protease recognition sequence. Fzd4 is additionally tagged with an N-terminal FLAG tag and Tspan12 is additionally tagged with a C-terminal 6xHis tag. The Fzd4 C-terminal PDZ ligand (ETVV) is appended after split GFP to improve expression and surface localization (Bruguera et al., 2022). **B.** Schematic of Tspan12/Fzd4 heterodimer expression and purification. The constructs in A. were co-expressed in Sf9 cells, solubilized in DDM, and co-purified on M1 anti-FLAG affinity resin followed by size exclusion chromatography. The dimer was reconstituted into nanodiscs and further purified by size exclusion followed by capture on anti-GFP nanobody resin, from which it was eluted by 3C protease. **C.** Size exclusion traces of empty nanodiscs or Tspan12/Fzd4 dimer reconstituted into excess nanodiscs, with absorbance detected at 280 nm and 475 nm (GFP absorption peak), on a Superose 6 Increase column. Indicated fractions were pooled for **D.** SDS-PAGE, imaged using StainFree imaging (above) or GFP fluorescence (below); intact GFP is SDS-resistant. Fractions on the right side of the peak (boxed) were pooled in order to exclude any potential separately-reconstituted dimers (i.e. two nanodiscs with one receptor each, linked together by the GFP moiety). **E.** Pooled fractions were purified by GFP nanobody resin and eluted with 3C protease. The load, flowthrough and eluate are shown.

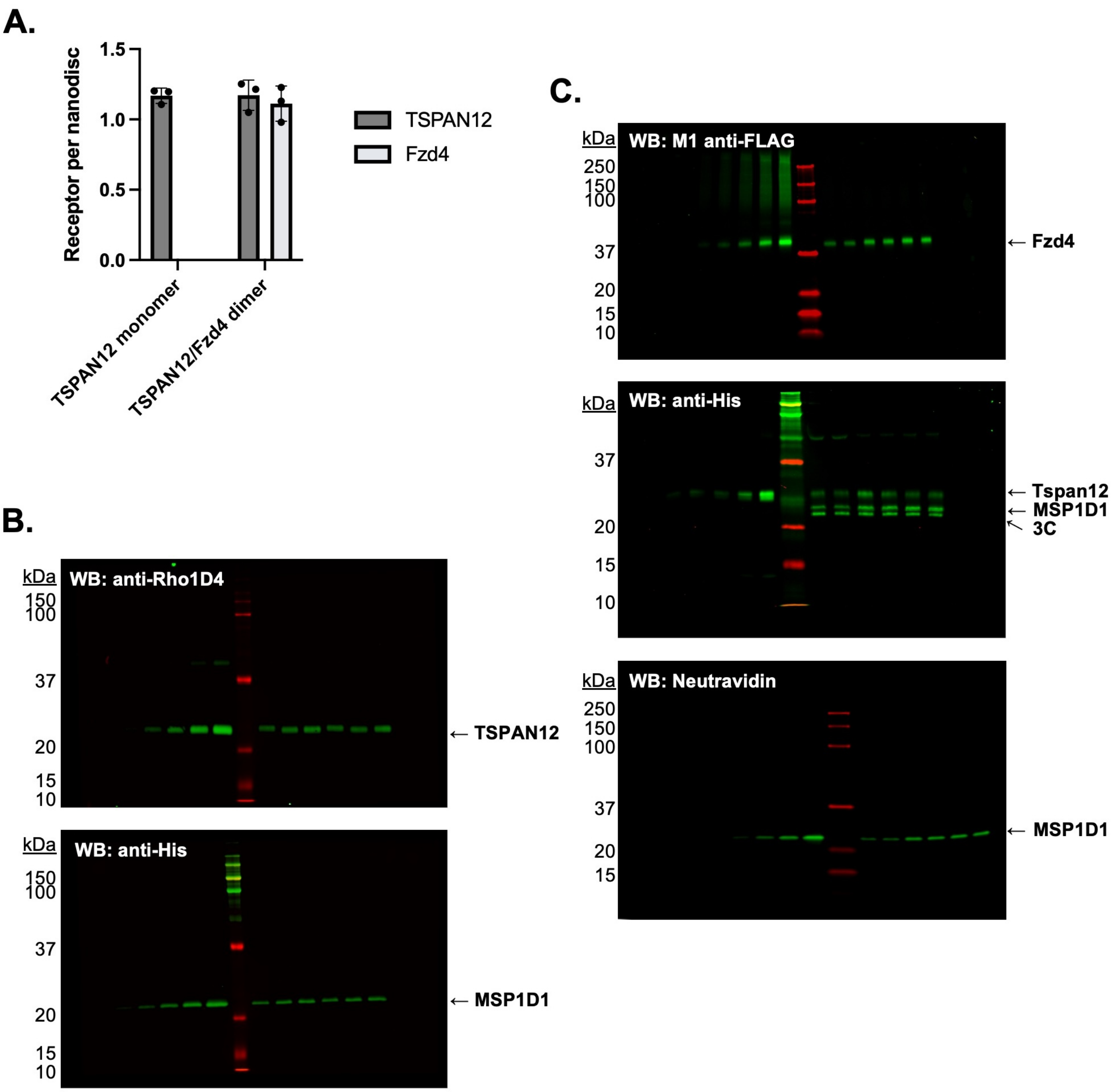


**Figure S9. Stoichiometry of receptors in nanodiscs was determined by quantitative western blot. A.** Receptors per nanodisc in monomeric Tspan12 (mean ± S.E.M. 1.17 ± 0.05 Tspan12 per two MSP1D1) and heterodimeric Tspan12/Fzd4 nanodiscs (1.17 ± 0.11 Tspan12 and 1.11 ± 0.12 Fzd4 per two MSP1D1) as calculated from three independent samples each, with each component measured three times each by quantitative western blot. **B.** Representative anti-Rho1D4 (top) and anti-his (bottom) western blots to quantify Tspan12-1D4 and His-MSP1D1, respectively, in monomeric Tspan12 reconstitutions. A known dilution series of purified protein was loaded in left lanes to generate a standard curve in the linear range of detection, against which dilutions of three nanodisc reconstitutions, loaded in duplicate in right lanes, were compared. **C.** Representative anti-FLAG (top), anti-His (middle) and Neutravidin-800 (bottom) western blots to quantify FLAG-Fzd4, Tspan12-His, and biotinylated MSP1D1, respectively, in Tspan12/Fzd4 heterodimer preparations.

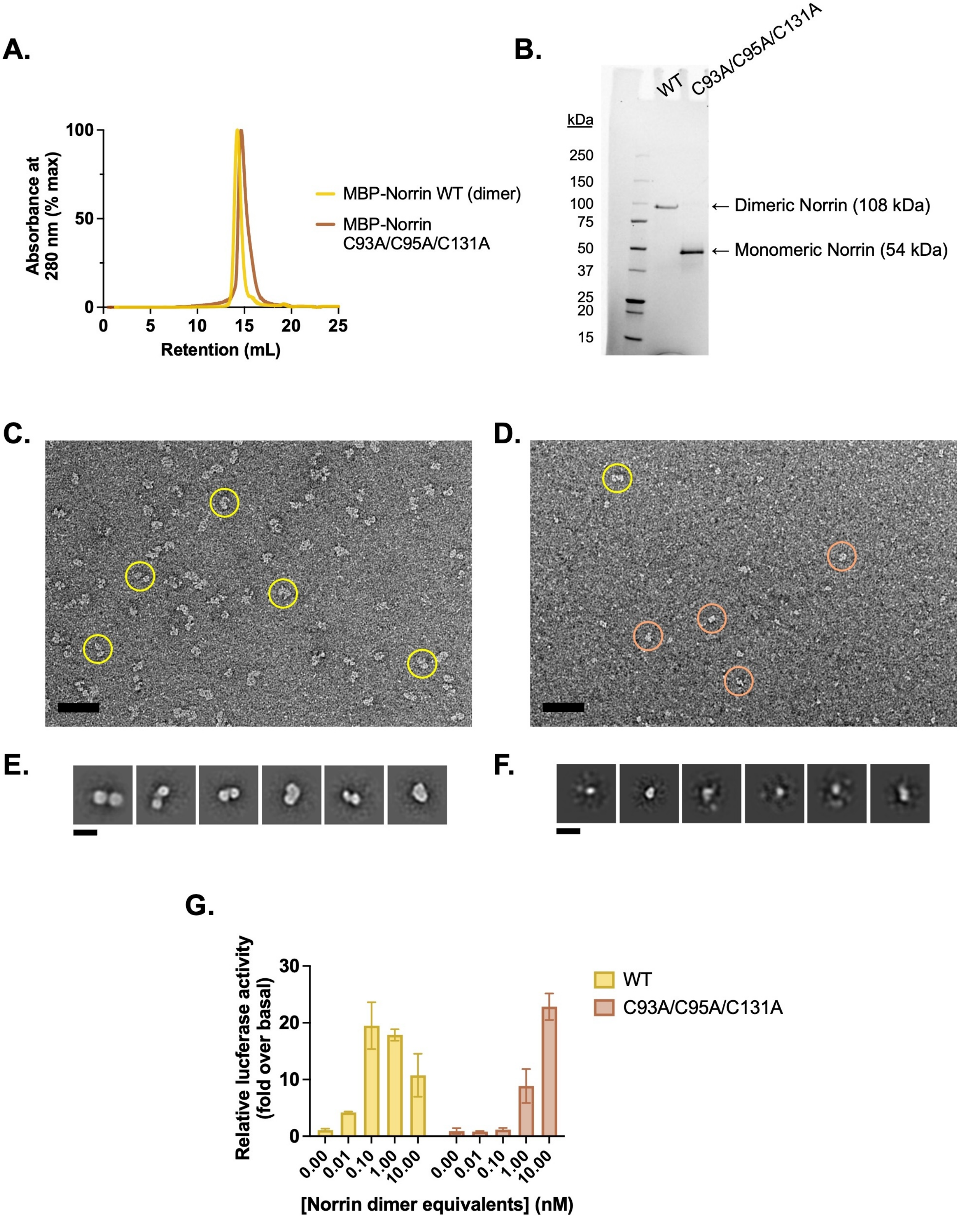


**Figure S10. Purification and validation of monomeric Norrin. A.** Analytical size exclusion traces of wildtype (dimeric) MBP-Norrin (yellow) and MBP-Norrin rendered monomeric (brown) via mutations C93A/C95A/C131A to eliminate the intermolecular disulfides. Purified protein was injected at 25 $\mu$M on a Superdex 200 Increase 10/300 column, resulting on an on-column concentration in excess of 2.5 $\mu$M assuming a 10-fold on-column dilution factor. **B.** Non-reducing SDS-PAGE gel of wildtype and C93A/C95A/C131A MBP-Norrin. **C.** Uranyl acetate negative stain micrograph of wildtype MBP-Norrin, prepared at 100 nM. Scale bar is 50 nm. Representative picked particles indicated in yellow. **D.** Uranyl acetate negative stain micrograph of MBP-Norrin C93A/C95A/C131A, prepared at 100 nM. Scale bar is 50 nm. Representative picked particles indicated with circles. Yellow-circled particle appears to be large enough to potentially be a dimer; brown circles show some smaller species, which dominate. **E.** 2D class averages of picked particles from C. show two lobes, consistent with two copies of MBP-Norrin (54 kDa each). **F.** 2D class averages of picked particles from D. show small, single particles that are hard to align; they are about half the size of particles in E, consistent with one copy of MBP-Norrin. This suggests that MBP-Norrin C93A/C95A/C131A is monomeric at 100 nM. **G.** β-catenin transcriptional activity in response to 0.01-10 nM purified WT (dimeric) or 0.02-20 nM C93A/C95A/C131A (monomeric) Norrin, in Fzd1/2/4/5/7/8-knockout HEK293T cells transfected with Fzd4 and TopFlash luciferase reporter plasmids. Data are plotted as mean ± S.D. from n=3 replicate wells.

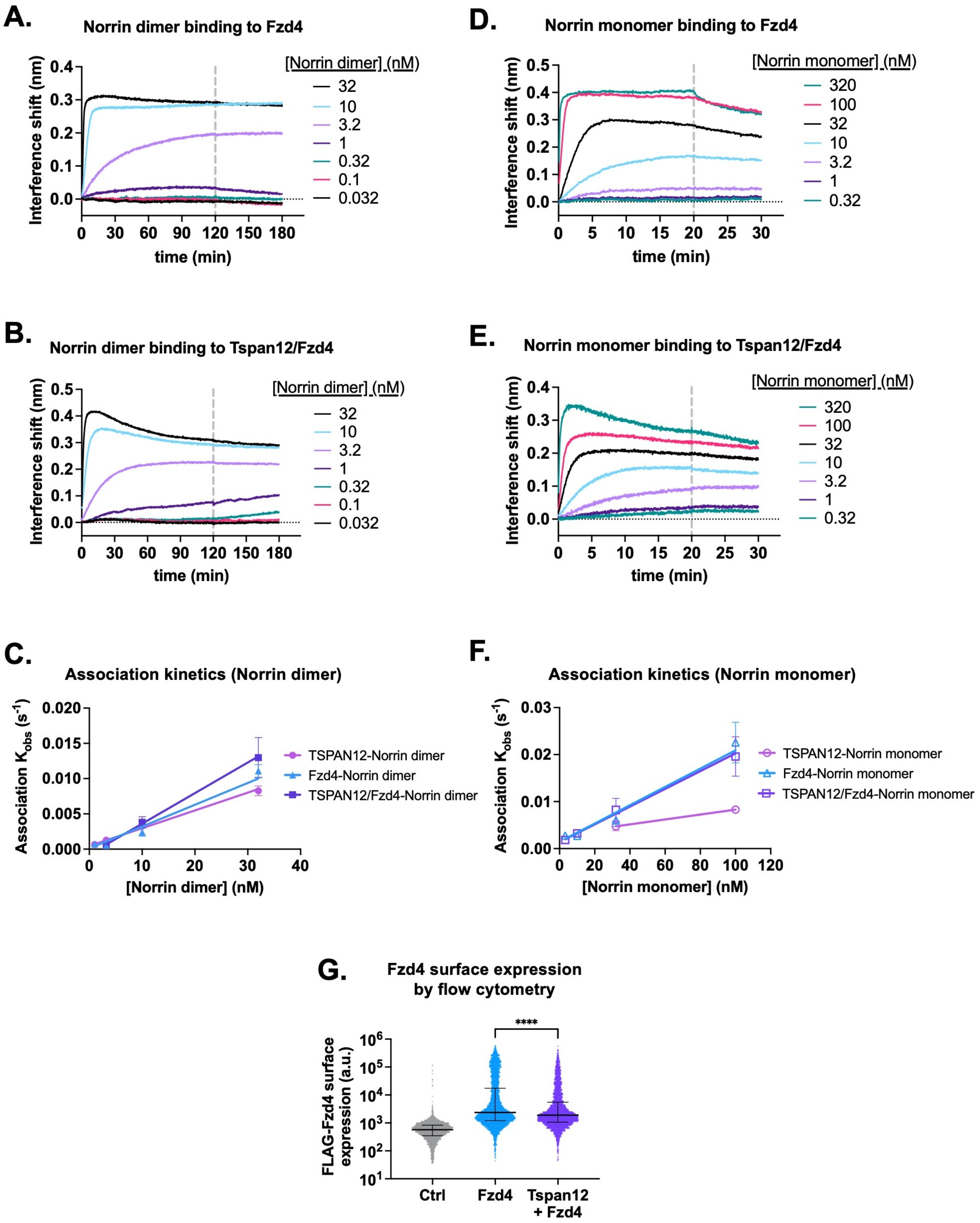


**Figure S11. Tspan12 enhances Norrin recruitment. A.** Representative BLI association and dissociation traces of dimeric Norrin binding to Fzd4 monomer or **B.** Tspan12/Fzd4 heterodimer in nanodiscs. **C.** Observed association rate constant $K_{obs}$ of Norrin dimer binding to Tspan12, Fzd4, or Tspan12/Fzd4 heterodimer in nanodiscs, plotted against Norrin concentration. Linear fits were used to obtained association rate constants reported in table S1. Data represent mean ± S.D. for three independent replicates**. D.** Representative BLI association and dissociation traces of monomeric Norrin (C93A/C95A/C131A) binding to Fzd4 monomer or **E.** Tspan12/Fzd4 heterodimer in nanodiscs. **F.** Observed association rate constant $K_{obs}$ (mean ± S.D.) of Norrin monomer binding to Tspan12, Fzd4, or Tspan12/Fzd4 heterodimer in nanodiscs, plotted against Norrin monomer concentration**. G.** Fzd4 surface expression on Expi293 cells transfected with empty vector,

FLAG-Fzd4, or FLAG-Fzd4 + Tspan12, which were then stained with M1 anti-FLAG antibody conjugated to Alexa Fluor 647. Cell fluorescence is measured by flow cytometry and plotted along with the median and interquartile range. Co-expression of Tspan12 modestly but significantly decreases surface expression of Fzd4 (Mann-Whitney test, p-value <0.0001 in each of three independent experiments).

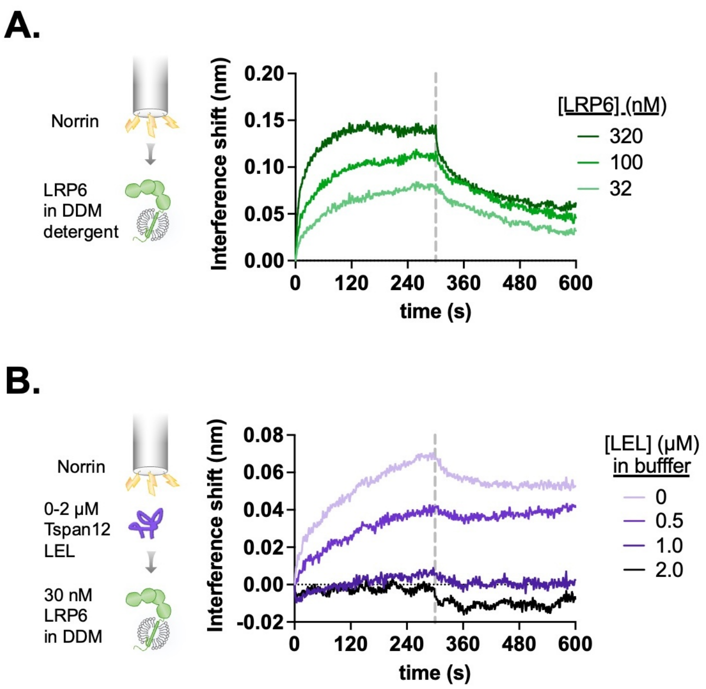


**Figure S12. LRP6-Norrin binding is reduced by Tspan12 LEL. A.** Association and dissociation biolayer interferometry traces of purified LRP6 (residues 20-1439, including the transmembrane domain but with a truncated C-terminus, in DDM detergent, 32, 100, or 320 nM) binding to MBP-Norrin-loaded biosensors. **B.** Biolayer interferometry traces of 30 nM LRP6 associating to and dissociating from MBP-Norrin-loaded biosensors, pre-equilibrated with increasing concentrations of MBP-fused Tspan12 LEL.
